# Novel *emm*4 lineage associated with an upsurge in invasive group A streptococcal disease in the Netherlands, 2022

**DOI:** 10.1101/2022.12.31.522331

**Authors:** Boas C.L. van der Putten, Wendy C.M. Bril-Keijzers, Lidewij W. Rümke, Stefan M.T. Vestjens, Linda A.M. Koster, Marloes Willemsen, Marlies A. van Houten, Nynke Y. Rots, Bart J.M. Vlaminckx, Brechje de Gier, Nina M. van Sorge

## Abstract

Invasive group A streptococcal (iGAS) disease cases increased in the first half year of 2022 in the Netherlands with a remarkably high proportion of *emm*4 isolates. Whole-genome sequence analysis of 66 *emm*4 isolates, 40 isolates from the pre-COVID-19-pandemic period 2009-2019 and 26 contemporary isolates from 2022, identified a novel *Streptococcus pyogenes* lineage (M4_NL22_), which accounted for 85% *emm*4 iGAS cases in 2022. Surprisingly, we detected few isolates of the *emm*4 hypervirulent clone, which has replaced nearly all other *emm*4 in the USA and the UK. M4_NL22_ displayed genetic differences compared to other *emm*4 strains, although these were of unclear biological significance. In publicly available data, we identified a single Norwegian isolate belonging to M4_NL22_, which was sampled after the isolates from this study, possibly suggesting export of M4_NL22_ to Norway. In conclusion, our study identified a novel *S. pyogenes emm*4 lineage underlying an increase of iGAS disease in early 2022 in the Netherlands and results have been promptly communicated with public health officials.

**Impact statement:** Group A *Streptococcus* (predominantly comprising *Streptococcus pyogenes*) has a long history of causing disease in humans. Over the past decades, researchers have described various changes in the *S. pyogenes* population circulating in humans. In this study, we describe a shift in the *S. pyogenes* population within type *emm*4 in the Netherlands in early 2022. A novel lineage, which we termed M4_NL22_, became the predominant *emm*4 lineage during this period. This was surprising, as in other countries such as the USA and the UK, another recently emerged lineage became dominant. Our work informed public health officials on the possible causes of increased *S. pyogenes* cases in the Netherlands and underlines the added value of continued active bacteriological surveillance.

**Data summary:** Code is freely available from https://github.com/boasvdp/GAS_M4. All sequencing data is freely available from BioProject PRJEB58654.

## Introduction

*Streptococcus pyogenes* can cause a wide variety of infections in humans, including non-invasive manifestations such as scarlet fever, pharyngitis and impetigo but also invasive manifestations such as fasciitis necroticans, streptococcal toxic shock syndrome, and puerperal fever. These three invasive group A streptococcal (iGAS) infections are notifiable by law in the Netherlands. An upsurge in both notifiable and non-notifiable iGAS infections was observed in the Netherlands compared to pre-COVID-19 levels in spring 2022, shortly after COVID-19-related restrictions were lifted. Alarmingly, this increase disproportionally affected children under 6 years of age^1,2^. Changes or shifts in dominance of *emm*-types, which are detected through bacteriological surveillance, may indicate changes in virulence or transmission within the *S. pyogenes* population.

Active bacteriological surveillance for *S. pyogenes* was initiated as part of a research project in 2019. *S. pyogenes* isolates cultured from normally-sterile compartments have been voluntarily submitted by medical microbiology laboratories to the Netherlands Reference Laboratory for Bacterial Meningitis (NRLBM) for *emm*-typing. Between 1 January and 13 May 2022, 25 isolates were received from children under 5 years of age of which *emm*4 and *emm*12 were the most dominant *emm* types (28% each), followed by *emm*1.0 (16%)^2^. Among isolates from all ages (n=134), *emm*4 ranked third (13%). The dominance of *emm*4 among iGAS cases was unexpected, since data from a retrospective study (2009-2019; L.W. Rümke, unpublished data) indicated that this *emm*-type was associated with carriage (7% of carriage isolates) rather than invasive disease (3% of invasive isolates) in the Netherlands.

Recently, a hypervirulent *emm*4 clone (henceforth referred to as the “hypervirulent clone”) was described, which has replaced nearly all other *emm*4 *S. pyogenes* strains in the USA and UK^3^. We hypothesized that this hypervirulent *emm*4 clone was responsible for the increase of *emm*4 iGAS cases in the Netherlands.

## Methods

### Sample selection

Since 2019, nine Dutch sentinel medical microbiology laboratories (MMLs; see Supplementary information for details) covering approximately 28% of the Netherlands, have been requested to submit *S. pyogenes* isolates for *emm*-typing to the Netherlands Reference Laboratory for Bacterial Meningitis (NRLBM) when cultured from normally-sterile compartments (including blood, cerebrospinal fluid [CSF], deep wounds, abscesses, et cetera). Isolates from the female reproductive tract (not a normally-sterile compartment) were associated with the clinical manifestation of puerperal fever/sepsis, a notifiable disease in the Netherlands, and were therefore counted as iGAS. This bacteriological surveillance was part of a research project to gain insight in the incidence of and molecular epidemiology of invasive group A streptococcal (iGAS) infections, beyond the iGAS notifiable manifestations (streptococcal toxic shock syndrome (STSS), puerperal fever and necrotizing fasciitis) in the Netherlands. From mid-April 2022, this request was expanded to all MMLs in the Netherlands. We included all 26 *emm*4 *S. pyogenes* isolates that were received as part of this active prospective surveillance between January 1^st^ and May 14^th^, 2022 (2022 cohort, Table 1). Additionally, 40 *emm*4 *S. pyogenes* isolates from the 2009-2019 period (2009-2019 or ‘pre-pandemic’ cohort) were included. These isolates were obtained from patients with invasive disease (n=11; Table 1) or from asymptomatic carriers (n=29; Table 1), originating from two studies. A total of 304 carriage isolates were obtained from the OKIDOKI study series (OKIDOKI 1-5), which are cross-sectional studies conducted triennially in the western part of the Netherlands since 2009 to assess the impact of vaccination on bacterial carriage^4^. Isolates were cultured from nasopharyngeal and oropharyngeal swabs of healthy 11-, 24- and 46-month old children and their parents. The 11 invasive *S. pyogenes* isolates were part of a retrospectively collected cohort of strains from the period 2009-2019. The total collection of this cohort consisted of 269 blood or CSF cultures from patients with iGAS disease that had been admitted to three Dutch hospitals in the middle of the Netherlands (Diakonessenhuis, Utrecht; University Medical Centre Utrecht, Utrecht; St. Antonius hospital, Nieuwegein). All 66 isolates were subjected to whole-genome sequencing (WGS).

**Table 1.** *S. pyogenes emm*4 isolate characteristics from the 2009-2019 and 2022 cohorts.

|  | 2009-2019 cohort |  | 2022 cohort |
| --- | --- | --- | --- |
|  | Carriage | Invasive | Invasive |
| Number of isolates | 29 | 11 | 26 |
| Median year of isolation (IQR) | 2012 (2009-2016) | 2013 (2013-2015) | 2022 |
| Cultured from adult*, no. (%) | 6 (21%) | 8 (72%) | 17 (65%) |
| Site of isolation |  |  |  |
| Blood, no. (%) | 0 (0%) | 11 (100%) | 15 (58%) |
| Nose, no. (%) | 23 (79%) | 0 (0%) | 0 (0%) |
| Throat, no. (%) | 6 (21%) | 0 (0%) | 0 (0%) |
| Female genital tract, no. (%) | 0 (0%) | 0 (0%) | 5 (19%) |
| Other<sup>#</sup>, no. (%) | 0 (0%) | 0 (0%) | 6 (23%) |
| Lineage |  |  |  |
| M4<sub>NL22</sub> lineage, no. (%) | 1 (3%) | 1 (3%) | 22 (85%) |
| Hypervirulent clone, no. (%) | 3 (10%) | 1 (9%) | 2 (8%) |

\*For one group, we only have age categories available instead of exact age
<sup>#</sup>Other sources include wound, abscess, general tissue and a dialysis machine

### Whole-genome sequencing

#### Short read sequencing

DNA extraction was performed using a magnetic bead-based protocol on an automated Maxwell machine (Promega). Whole-genome sequencing of 62 (of total 66) isolates was conducted by the Core Facility Genomics of the Amsterdam UMC. Library preparation was performed using the Kapa HTP Library Preparation kit and libraries were sequenced on an Illumina Hiseq4000 using a PE150 kit, all according manufacturer’s instructions. The remaining four isolates were sequenced by MicrobesNG, United Kingdom (https://microbesng.com/). Isolates were suspended in DNA Shield buffer (Zymo Research, USA) and shipped to MicrobesNG. The DNA extraction, library preparation, sequencing and bioinformatics QC protocols of MicrobesNG are available online (v20210419, https://microbesng.com/documents/24/MicrobesNG_Sequencing_Service_Methods_v20210419.pdf).

#### Long read sequencing

Isolate 2220758 was selected for long read sequencing using Oxford Nanopore Technologies (ONT). Reads from three sequencing experiments were combined due to low yields of individual experiments. DNA extractions for the first two sequencing experiments were performed according to a method described previously^5^. DNA extraction for the third sequencing experiment was performed using the Maxwell RSC Cultured Cells DNA Kit (Cat.# AS1620) according to manufacturer’s instructions. Library preparation and MinION sequencing methodologies for all three experiments were described before^5^.

### WGS analysis

#### Short read analysis

Illumina reads were processed with an adapted version of the NRLBM genomic pipeline (v0.3, https://github.com/NRLBM/assembly/releases/tag/v0.3). All analyses were run with default settings unless otherwise noted. Sequence reads were trimmed and filtered using Trimmomatic v0.36^6^. Sequence read qualities were assessed before and after Trimmomatic using FastQC v0.11.9^7^. Contamination was assessed using Kraken2 v2.1.2^8^ and the MiniKraken v1 database. Filtered sequence reads were assembled using the Shovill v1.1.0 wrapper (https://github.com/tseemann/shovill) for SPAdes v3.15.4^9^. Depth of coverage was assessed by mapping filtered reads on de novo assemblies using minimap2 v2.17-r941^10^, samtools v1.9^11^ and bedtools v2.29.0^12^. Assembly quality was assessed using Quast v4.6.3^13^. Multi-locus sequence types were assigned using the PubMLST API^14^. Virulence genes were detected using ABRicate v1.0.1 (https://github.com/tseemann/abricate) and VFDB^15^. *emm*-types were confirmed using emmtyper v0.2.0 (https://github.com/MDU-PHL/emmtyper). Antimicrobial resistance genes and mutations were identified using AMRfinderplus, with curated thresholds for *S. pyogenes* (“--organism Streptococcus_pyogenes”)^16^. Domains of individual protein sequences were investigated using the InterPro webserver^17^.

A maximum-likelihood (ML) phylogeny was constructed by mapping filtered sequence reads on the Duke strain reference genome (GenBank accession number CP031770) using Snippy v4.6.0 (https://github.com/tseemann/snippy). The previously described hypervirulent clone was identified according to 36 SNPs typical for this lineage^3^. A genome alignment was created from Snippy mappings and recombination was identified using ClonalFrameML v1.12^18^. Recombinatory regions were masked using maskrc-svg v0.5 (https://github.com/kwongj/maskrc-svg) and a ML phylogeny was inferred from the recombination-free alignment using IQ-TREE v2.0.3^19^, with the substitution model HKY+F.

Draft assemblies were annotated using Bakta v1.4.1^20^ with database v3.1. The pan-genome of this dataset was defined using Panaroo v1.2.10^21^ in strict mode and lineage-associated genetic elements were identified using PySEER v1.3.9^22^. A Bonferroni correction was applied to define statistically significant associations.

#### Long read analysis

Leading and trailing 80 bp of all ONT reads were trimmed using NanoFilt v2.8.0^23^. Trimmed reads were subsequently filtered using filtlong v0.2.1 (https://github.com/rrwick/Filtlong), retaining reads of at least 5,000 bp long and the best 90% of reads matched by nucleotide identity based on Illumina reads. Illumina and ONT reads were assembled using Unicycler v0.5.0^24^ resulting in a completely resolved genome. The complete genome of 2220758 was annotated as described above. Additionally, the PHASTER webserver^25^ was queried to annotate prophages and the Artemis Comparison Tool v18.2.0^26^ was used to identify large-scale genomic rearrangements. The ICEfinder webserver was used to identify Integrative Conjugative Elements (ICEs)^27^.

#### Public data

To relate the Dutch data to other recent surveillance data of *S. pyogenes*, we downloaded reads from BioProjects PRJNA395240 and PRJEB42599 comprising *S. pyogenes* data from USA and Norwegian surveillance programmes, respectively. Downloaded reads were processed together with Dutch data using the standard pipeline. We could not identify other sources of publicly available real time genomic surveillance of *S. pyogenes*.

## Results

### Patient characteristics

For the 26 isolates from the 2022 cohort, information on age and sex was available for 25 isolates. Median patient age was 34 years (IQR: 5-43 years) and 15 (60%) were male. Site of isolation is summarized in Table 1. Of the 2009-2019 cohort, 26 out of 40 (65%) were from 0-5-year olds, while the remaining 14 isolates were from adults.

### Bacterial characteristics: new *emm*4 clone

The hypervirulent clone represented less than 10% among recent and older *S. pyogenes* isolates (Table 1), contrasting the strong emergence of this clone in the USA and the UK. Moreover, 22 out-of-26 (85%) *emm*4 iGAS isolates from the 2022 cohort clustered together, representing a new *emm*4 lineage M4_NL22_ (Figure 1), whereas only two out-of-40 (5%) isolates from the 2009-2019 cohort (isolated in 2017 and 2018) belonged to this M4_NL22_ lineage. Fifteen out-of-22 (68%) M4_NL22_ isolates showed very limited genetic variation with a median of 6 SNPs between them (range 0-16), suggesting a recent emergence and potentially a high transmission rate. Indeed, analysis of 46 isolates obtained from a recent cross-sectional carriage study (Sept 2022-Feb 2023; OKIDOKI-6) among 24-48-month-old children and their parents, *emm*4 was the dominant *emm*-type (n=13, 28%). Unfortunately, no WGS data are currently available to confirm whether these *emm*4.0 isolates belong to the new M4_NL22_ lineage.

**Figure 1.**
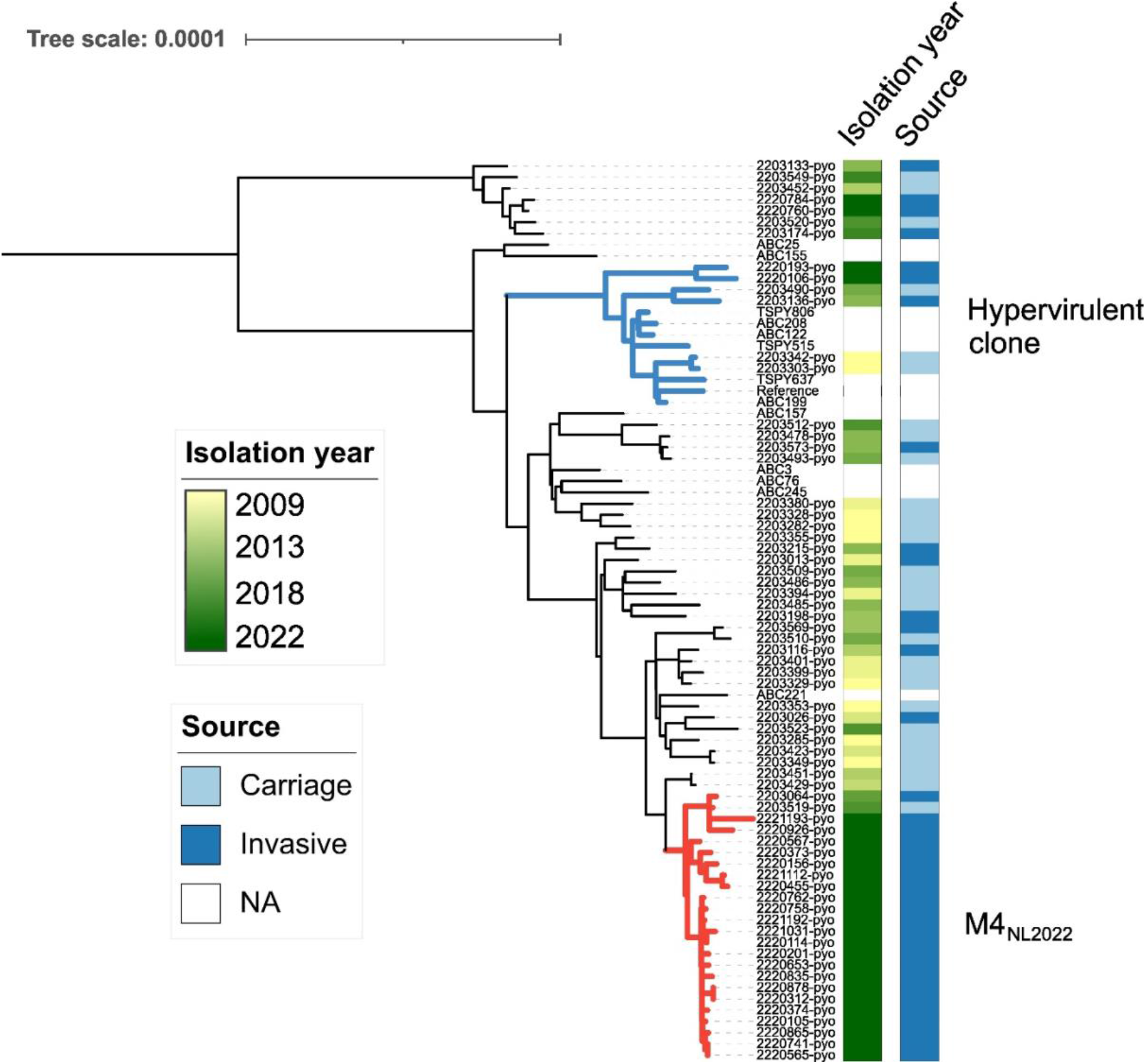
Phylogeny of 66 Dutch *emm*4 isolates and 14 reference genomes. Reference genomes originate from Debroy *et al*. (2021) (*2*) and are shown without metadata. The hypervirulent clone lineage is colored blue and the M4_NL22_ lineage is colored red. An interactive version of these data with metadata is available at https://itol.embl.de/tree/92108202133469351670785481#.

Analysis of public surveillance databases, identified one isolate (NCBI BioSample SAMEA111521236) from 2022 from Norway (Norwegian Public Institute of Health surveillance, NCBI BioProject PRJEB42599) that clustered in the clonal part of the M4_NL22_ lineage and differed 9 SNPs from the closest Dutch isolate. No M4_NL22_ isolates were identified from USA surveillance data from 2021-2022 (CDC Active Bacterial Core surveillance, NCBI BioProject PRJNA395240). No resistance genes or mutations were detected in any Dutch isolate.

### Genetic characteristics of M4_NL22_ lineage strains

The M4_NL22_ lineage displayed several genetic variations compared to other *S. pyogenes emm*4 strains, including six unique SNPs and a 306 bp deletion in the *proA* gene, which is involved in proline biosynthesis (see Table S2 in the Appendix for details). The 306 bp deletion in *proA* likely results in a defective protein, since it deletes almost a quarter of the entire predicted protein length (amino acids 129-230), which overlaps with an aldehyde dehydrogenase family domain (Pfam accession PF00171). Characteristically, M4_NL22_ lacked the *hasABC* genes for synthesis of a hyaluronic acid capsule^28^. Further characterisation through *in vitro* and *in vivo* experiments should reveal whether M4_NL22_ is more transmissible than other *emm*4 *S. pyogenes*. No genetic changes in known virulence determinants were observed (*nga*-*ifs*-*slo* region, superantigens, known virulence genes) in isolates from this M4_NL22_ lineage.

The complete genome of isolate 2220758 comprised 1,890,346 bp, which is approximately 13.5 kbp smaller than the Duke *emm*4 reference genome. Based on pairwise blastn analysis, 1817 genes were shared between Duke and 2220758 (92.7% of all 1960 genes of 2220758). Of the 143 genes uniquely present in 2220758 compared to the Duke reference genome, 23 (16%) were annotated as phage protein-coding genes. The remaining genes were annotated with a variety of functions.

The complete genome of isolate 2220758 harboured two intact prophages (closest PHASTER hits: Streptococcus phage 315.3, NCBI Nucleotide accession NC_004586, and Streptococcus phage 315.5, NC_004588), one possibly intact prophage (closest hit: Streptococcus phage 315.2, NC_004585, marked “questionable” by PHASTER) and two incomplete prophages. Streptococcus phage 315.3 harbours virulence-associated genes *hylP* and *ssa*, while Streptococcus phage 315.5 harbours *fbp54* and *mf3*. In comparison to isolate ABC221, representing the genome most closely related to M4_NL22_, isolate 2220758 did not harbour Streptococcus phage 315.4, which typically encodes a streptococcal DNase *sdn*. However, 14 of the M4_NL22_ isolates did harbour the *sdn* gene, and also fragments of Streptococcus phage 315.4 (mean: 53% coverage). Although phages are challenging to detect in fragmented draft assemblies, all *emm*4 isolates harboured at least some fragment of Streptococcus phage 315.3 (mean: 79.9% coverage) and Streptococcus phage 315.5 (mean: 74.7% coverage). Twenty-three out-of-24 (96%) M4_NL22_ isolates also harboured fragments of Streptococcus phage 315.2, albeit with lower coverage (mean: 45%). Isolate 2220758 also harboured one putative Integrative and Mobilizable Element (IME), containing *grab*, a gene encoding an alpha2-macroglobulin-binding protein^29^, and one putative Integrative Conjugative Element (ICE), harbouring *znuBC-mtsA*, which possibly encodes an ABC transporter for manganese uptake. Finally, this ICE also harboured small non-coding RNA Spy490380c (Rfam accession RF02644) present in multiple streptococcal species.

## Discussion

Several novel *S. pyogenes* clones have emerged over the past years, including *emm*4^3^, *emm*89^30^, and *emm*1 lineages^31^. Although most emerged lineages seem endowed with increased capacity for transmission or invasiveness, there may be additional explanations for the emergence of M4_NL22_ reported in the current study. As with many other infectious diseases^32^, we have observed a marked shift in the epidemiology of *S. pyogenes* infections after COVID-19 pandemic restrictions were lifted, with a surprising absolute and proportional increase of *emm*4 isolates among iGAS cases^2^.

There were only a few genetic differences between M4_NL22_ and other *emm4 S. pyogenes*, including six SNPs and a unique in-frame deletion of *proA*, a gene involved in proline biosynthesis. The biological implications of these six SNPs and the *proA* deletion are unclear and should be investigated in more detail. No large-scale genomic rearrangements were observed for M4_NL22_. However, 54% of M4_NL22_ isolates harboured the streptococcal DNase *sdn*, encoded by Streptococcus phage 315.4. Interestingly, the lack of this prophage in the complete genome of 2220758 may suggest that the phage is able to move through the *S. pyogenes emm*4 population within a short time span. Remmington *et al*. previously showed that the hypervirulent clone of *emm*4 *S. pyogenes* contained many degraded and probably immobile prophages^33^. However, prophages in M4_NL22_ did not seem to be degraded or immobile based on our sequence analysis, which is expected as M4NL22 has not descended from the hypervirulent clone.

Our study has several limitations. First, our sample size is relatively small and lacks data from active routine national surveillance for the period before 2019. Second, our study lacks phenotypic characterization of the novel M4_NL22_ lineage. Even though epidemiology suggests this lineage is well-adapted to spread and cause disease, this should be tested experimentally. Finally, although M4_NL22_ seems to have spread successfully in the Netherlands, only a single isolate of this lineage has been identified abroad. Although this may suggest that M4_NL22_ is mainly restricted to the Netherlands, another plausible explanation is the scarcity of real-time, publicly available genomic surveillance data of *S. pyogenes*, limiting comparisons to these countries.

In conclusion, we identified an emerging *emm*4 lineage in the Netherlands. Based on the dominance of *emm*4 among asymptomatic carriers of *S. pyogenes* between September 22 and February 2023, it is likely that this lineage descended from a predominantly carriage-associated type. Although iGAS cases in late 2022 and early 2023 in the Netherlands are dominated by *emm*1 (more specifically M1UK)^2^, *emm*4 isolates are continuously detected in patients with invasive disease. Continued bacteriological surveillance of isolates from both invasive disease and asymptomatic carriers remains necessary to timely inform public health officials of circulating clones with possibly altered transmission or virulence capacities. Genomic analysis is required to do this effectively, as classical typing techniques (e.g. *emm*-typing), while valuable, have insufficient resolution to provide valuable information for strain comparison.

## Acknowledgments

The authors would like to thank Agaath Arends-van ‘t Klooster, Ilse de Beer-Schuurman and Claudia Burger for their continued contributions as part of the NRLBM as well as the medical microbiology laboratories for their continued submission of isolates.

We thank SURF (www.surf.nl) for the support in using the Lisa Compute Cluster.

We thank Jeroen Bos, Sietze Brandes, Fabian Landman, Sjoerd Kuiling and Rob Mariman of the National Institute for Public Health and the Environment (RIVM) for the ONT sequencing of strain 2220758.

## Appendix

- Supplementary information.

- Table S1: Isolates included in this study.

- Table S2: Annotations of SNPs associated with M4_NL22_

## References

1. van Kempen EB, Bruijning-Verhagen PCJ, Borensztajn D, et al. Increase in invasive group A streptococcal infections in children in the Netherlands, a survey among 7 hospitals in 2022. Pediatr Infect Dis J. 2023;42(4):e122–e124. doi:10.1097/INF.0000000000003810

2. Gier B de, Marchal N, Beer-Schuurman I de, et al. Increase in invasive group A streptococcal (*Streptococcus pyogenes*) infections (iGAS) in young children in the Netherlands, 2022. Eurosurveillance. 2023;28(1):2200941. doi:10.2807/1560-7917.ES.2023.28.1.2200941

3. DebRoy S, Sanson M, Shah B, et al. Population Genomics of *emm*4 Group A *Streptococcus* Reveals Progressive Replacement with a Hypervirulent Clone in North America. mSystems. 2021;6(4):e00495–21. doi:10.1128/mSystems.00495-21

4. Bosch AATM, van Houten MA, Bruin JP, et al. Nasopharyngeal carriage of *Streptococcus pneumoniae* and other bacteria in the 7th year after implementation of the pneumococcal conjugate vaccine in the Netherlands. Vaccine. 2016;34(4):531–539. doi:10.1016/j.vaccine.2015.11.060

5. Hendrickx APA, Landman F, de Haan A, et al. Plasmid diversity among genetically related *Klebsiella pneumoniae* blaKPC-2 and blaKPC-3 isolates collected in the Dutch national surveillance. Sci Rep. 2020;10:16778. doi:10.1038/s41598-020-73440-2

6. Bolger AM, Lohse M, Usadel B. Trimmomatic: a flexible trimmer for Illumina sequence data. Bioinformatics. 2014;30(15):2114–2120. doi:10.1093/bioinformatics/btu170

7. Andrews S. FastQC a quality control tool for high throughput sequence data. URL: https://www.bioinformatics.babraham.ac.uk/projects/fastqc. Published online 2010.

8. Wood DE, Lu J, Langmead B. Improved metagenomic analysis with Kraken 2. Genome Biol. 2019;20(1):257. doi:10.1186/s13059-019-1891-0

9. Prjibelski A, Antipov D, Meleshko D, Lapidus A, Korobeynikov A. Using SPAdes De Novo Assembler. Current Protocols in Bioinformatics. 2020;70(1):e102. doi:10.1002/cpbi.102

10. Li H. Minimap2: pairwise alignment for nucleotide sequences. Bioinformatics. 2018;34(18):3094–3100. doi:10.1093/bioinformatics/bty191

11. Li H, Handsaker B, Wysoker A, et al. The Sequence Alignment/Map format and SAMtools. Bioinformatics. 2009;25(16):2078–2079. doi:10.1093/bioinformatics/btp352

12. Quinlan AR, Hall IM. BEDTools: a flexible suite of utilities for comparing genomic features. Bioinformatics. 2010;26(6):841–842. doi:10.1093/bioinformatics/btq033

13. Gurevich A, Saveliev V, Vyahhi N, Tesler G. QUAST: quality assessment tool for genome assemblies. Bioinformatics. 2013;29(8):1072–1075. doi:10.1093/bioinformatics/btt086

14. Jolley KA, Bray JE, Maiden MCJ. A RESTful application programming interface for the PubMLST molecular typing and genome databases. Database. 2017;2017:bax060. doi:10.1093/database/bax060

15. Liu B, Zheng D, Jin Q, Chen L, Yang J. VFDB 2019: a comparative pathogenomic platform with an interactive web interface. Nucleic Acids Research. 2019;47(D1):D687–D692. doi:10.1093/nar/gky1080

16. Feldgarden M, Brover V, Haft DH, et al. Validating the AMRFinder Tool and Resistance Gene Database by Using Antimicrobial Resistance Genotype-Phenotype Correlations in a Collection of Isolates. Antimicrob Agents Chemother. 2019;63(11). doi:10.1128/AAC.00483-19

17. Paysan-Lafosse T, Blum M, Chuguransky S, et al. InterPro in 2022. Nucleic Acids Research. 2023;51(D1):D418–D427. doi:10.1093/nar/gkac993

18. Didelot X, Wilson DJ. ClonalFrameML: Efficient Inference of Recombination in Whole Bacterial Genomes. PLOS Computational Biology. 2015;11(2):e1004041. doi:10.1371/journal.pcbi.1004041

19. Minh BQ, Schmidt HA, Chernomor O, et al. IQ-TREE 2: New Models and Efficient Methods for Phylogenetic Inference in the Genomic Era. Molecular Biology and Evolution. 2020;37(5):1530–1534. doi:10.1093/molbev/msaa015

20. Schwengers O, Jelonek L, Dieckmann MA, Beyvers S, Blom J, Goesmann A. Bakta: rapid and standardized annotation of bacterial genomes via alignment-free sequence identification. Microb Genom. 2021;7(11):000685. doi:10.1099/mgen.0.000685

21. Tonkin-Hill G, MacAlasdair N, Ruis C, et al. Producing polished prokaryotic pangenomes with the Panaroo pipeline. Genome Biology. 2020;21(1):180. doi:10.1186/s13059-020-02090-4

22. Lees JA, Croucher NJ, Goldblatt D, et al. Genome-wide identification of lineage and locus specific variation associated with pneumococcal carriage duration. Cobey S, ed. eLife. 2017;6:e26255. doi:10.7554/eLife.26255

23. De Coster W, D’Hert S, Schultz DT, Cruts M, Van Broeckhoven C. NanoPack: visualizing and processing long-read sequencing data. Bioinformatics. 2018;34(15):2666–2669. doi:10.1093/bioinformatics/bty149

24. Wick RR, Judd LM, Gorrie CL, Holt KE. Unicycler: Resolving bacterial genome assemblies from short and long sequencing reads. PLOS Computational Biology. 2017;13(6):e1005595. doi:10.1371/journal.pcbi.1005595

25. Arndt D, Grant JR, Marcu A, et al. PHASTER: a better, faster version of the PHAST phage search tool. Nucleic Acids Res. 2016;44(W1):W16–21. doi:10.1093/nar/gkw387

26. Carver TJ, Rutherford KM, Berriman M, Rajandream MA, Barrell BG, Parkhill J. ACT: the Artemis comparison tool. Bioinformatics. 2005;21(16):3422–3423. doi:10.1093/bioinformatics/bti553

27. Liu M, Li X, Xie Y, et al. ICEberg 2.0: an updated database of bacterial integrative and conjugative elements. Nucleic Acids Research. 2019;47(D1):D660–D665. doi:10.1093/nar/gky1123

28. Flores AR, Chase McNeil J, Shah B, Van Beneden C, Shelburne SA. III. Capsule-Negative emm Types Are an Increasing Cause of Pediatric Group A Streptococcal Infections at a Large Pediatric Hospital in Texas. Journal of the Pediatric Infectious Diseases Society. 2019;8(3):244–250. doi:10.1093/jpids/piy053

29. Toppel AW, Rasmussen M, Rohde M, Medina E, Chhatwal GS. Contribution of protein G-related alpha2-macroglobulin-binding protein to bacterial virulence in a mouse skin model of group A streptococcal infection. J Infect Dis. 2003;187(11):1694–1703. doi:10.1086/375029

30. Turner CE, Abbott J, Lamagni T, et al. Emergence of a New Highly Successful Acapsular Group A Streptococcus Clade of Genotype emm89 in the United Kingdom. mBio. 2015;6(4):e00622–15. doi:10.1128/mBio.00622-15

31. Lynskey NN, Jauneikaite E, Li HK, et al. Emergence of dominant toxigenic M1T1 *Streptococcus pyogenes* clone during increased scarlet fever activity in England: a population-based molecular epidemiological study. The Lancet Infectious Diseases. 2019;19(11):1209–1218. doi:10.1016/S1473-3099(19)30446-3

32. Chow EJ, Uyeki TM, Chu HY. The effects of the COVID-19 pandemic on community respiratory virus activity. Nat Rev Microbiol. Published online October 17, 2022:1–16. doi:10.1038/s41579-022-00807-9

33. Remmington A, Haywood S, Edgar J, Green LR, de Silva T, Turner CE. Cryptic prophages within a *Streptococcus pyogenes* genotype emm4 lineage. Microb Genom. 2020;7(1):mgen000482. doi:10.1099/mgen.0.000482

