## Supplemental information for "Novel *emm*4 lineage associated with an upsurge in invasive group A streptococcal disease in the Netherlands, 2022"

**Supplementary information**

### Information of medical microbiology laboratories that submitted *S. pyogenes* isolates

Nine sentinel laboratories submitting *S. pyogenes* isolates to the NRLBM since 2019:

Amsterdam UMC, Amsterdam; Rijnstate Hospital, Arnhem/Velp; Regionaal Laboratorium Medische Microbiologie Dordrecht/Gorinchem; Streeklaboratorium voor de Volksgezondheid, Haarlem; LabMicTa, Hengelo; Certe Medische Microbiologie Friesland & Noordoostpolder, Leeuwarden; St. Antonius Hospital, Nieuwegein; Zuyderland Medisch Centrum, Sittard; Eurofins PAMM, Veldhoven.

Other microbiology laboratories that submitted isolates for this study:

Erasmus UMC, Rotterdam; UMC Utrecht, Utrecht; OLVG, Amsterdam; Meander Medisch Centrum, Amersfoort; Reinier Haga Hospital, Delft; Diakonessenhuis Utrecht/Zeist/Doorn, Utrecht; Eurofins Clinical Diagnostics, Leiden; Regionaal Laboratorium, Den Bosch; MICROVIDA, Roosendaal.
